# Depletion of Polypyrimidine tract binding protein 1 (*ptbp1*) activates Müller glia-derived proliferation during zebrafish retina regeneration via modulation of the senescence secretome

**DOI:** 10.1101/2025.03.24.645057

**Authors:** Gregory J. Konar, Audrey L. Lingan, Kyle T. Vallone, Tu D. Nguyen, Zachary R. Flickinger, James G. Patton

**Author notes:** **Corresponding author:** James G Patton, PhD, Department of Biological Sciences, Vanderbilt University. Authors contributed equally to this work (Gregory J. Konar, Audrey L. Lingan).

## Abstract

Polypyrimidine Tract Binding protein 1 (PTB) is an alternative splicing factor linked to neuronal induction and maturation. Previously, knockdown experiments supported a model in which PTB can function as a potent reprogramming factor, able to elicit direct glia-to-neuron conversion *in vivo*, in both the brain and retina. However, later lineage tracing and genetic knockouts of PTB did not support direct neuronal reprogramming. Nevertheless, consistent with the PTB depletion experiments, we show that antisense knockdown of PTB (*ptbp1a*) in the zebrafish retina can activate Müller glia-derived proliferation and that depletion of PTB can further enhance proliferation when combined with acute NMDA damage. The effects of PTB are consistent with a role in controlling key senescence and pro-inflammatory genes that are part of the senescence secretome that initiates retina regeneration.

## Introduction

Degenerative eye diseases continue to be a leading cause of blindness, affecting millions of people worldwide. Conditions such as age-related macular degeneration (AMD) and retinitis pigmentosa target the retina, the innermost part of the eye composed of multiple layers of light-sensitive nervous tissue that project sensory information to the optic nerve. Damage or loss of one or more layers of the retina is irreversible in humans and other mammals. In contrast, zebrafish (*Danio rerio*) possess the ability to spontaneously regenerate lost neurons and cells of the retina in a Müller glia (MG)-dependent manner (Bernardos et al., 2007; Nagashima et al., 2013; Raymond et al., 2006; Yurco & Cameron, 2005). In response to damage zebrafish, MG undergo dedifferentiation and asymmetric division for self-replacement and the generation of a pool of proliferating neuronal progenitor cells that can replace all lost cell types (Fausett & Goldman, 2006; Raymond et al., 2006; Wan & Goldman, 2016; Yurco & Cameron, 2005). Extensive work is ongoing to identify genes and pathways that regulate zebrafish retina regeneration and determine why regeneration is blocked in humans (Lahne et al., 2020; Lyu et al., 2023; Nagashima & Hitchcock, 2021; Pavlou & Reh, 2023; Wan & Goldman, 2016). Immediately after damage, inflammation is activated, but as regeneration proceeds, there is a dynamic response that shifts from a pro-inflammatory state toward a pro-regenerative state (Bludau et al., 2024; Iribarne, 2021; Iribarne & Hyde, 2022b; Mitchell et al., 2018; Nagashima & Hitchcock, 2021; Silva et al., 2020). The inability to effect this switch leads to chronic inflammation, fibrosis and scarring (Iribarne & Hyde, 2022a; Mitchell et al., 2019; Palazzo et al., 2022; Todd et al., 2020; Todd et al., 2019).

In the retina, senescent cells play a key role in modulating inflammatory responses during regeneration through the release of factors that collectively make up the Senescence Associated Secretory Phenotype (SASP)(Coppé et al., 2010; He & Sharpless, 2017; Konar et al., 2024). Early after retina damage in zebrafish, a subset of macrophages and glial cells express markers of senescence and the release of SASP factors from these cells can modulate the microenvironment (Konar et al., 2024; Oubaha et al., 2016). As regeneration proceeds, senescent cells are cleared, coincident with a shift toward an anti-inflammatory, pro-regenerative response (Konar et al., 2024). Premature removal of senescent cells inhibits regeneration by altering the required dynamic changes in inflammation (Konar et al., 2024).

Recently, two different strategies were used to down-regulate the alternative splicing factor PTB (Van Nostrand et al., 2020) in the retina and the brain that suggested direct glia to neuron conversion with rescue of visual function after retina damage and replacement of dopaminergic neurons in a mouse model of Parkinson’s disease (Qian et al., 2020; Zhou et al., 2020). However, follow up lineage tracing and genetic knockout experiments did not support a role for PTB in direct neuronal conversion (Hoang et al., 2023; Wang et al., 2021; Xie et al., 2022). The conflicting results could be due to different experimental strategies, but might also be due to differences between knockdowns, knockouts, and genetic compensation or transcriptional adaptation (El-Brolosy & Stainier, 2017; Fu & Mobley, 2023; Rossi et al., 2015). Despite the controversy, we show here that antisense knockdown of PTB in otherwise undamaged zebrafish retinas leads to the generation of proliferating cells and further enhances the number of MG-derived proliferating cells when combined with NMDA damage. Consistent with a role for PTB in promoting senescence and regulating SASP expression (Georgilis et al., 2018; Hensel, Nicholas, Kimble, Nagpal, Omar, Tyburski, Jellison, Ménoret, et al., 2022; Li et al., 2024; Yang et al., 2024), depletion of PTB in the retina causes an increase in the number of senescent cells and expression of senescent markers. Together, our data argue that depletion of PTB contributes to retina regeneration by modulating the retinal microenvironment that facilitates MG-derived regeneration.

## Methods

### Zebrafish husbandry and maintenance

Experiments were performed using randomly selected male and female zebrafish, either wild-type AB or *Tg(1016tuba1a:GFP)* zebrafish. Zebrafish were kept at 28 °C on a 12:12 hour light-dark cycle. Experiments were conducted in accordance with Vanderbilt University Institutional Animal Care and Use Committee (IACUC) approval #M1800200.

### Antisense Oligonucleotide (ASO) Treatment

Antisense oligonucleotides (ASOs) were generated targeting *ptbp1a* using the following sequences:

**Zebrafish PTBP1a ASO 2:** 5’-TCTCTAAACGGCCGTCCATG-3’

**Zebrafish PTBP1a ASO 3:** 5’-GACGTAACTGGATCCCAGAGG-3’

**Zebrafish PTBP1a ASO 4:** 5’-GTGCAACCTGAAAGAGTGCG-3’

ASOs were synthesized by Sigma. All internucleotide linkages were phosphorothioates interspersed with phosphodiesters, and all cytosine residues were 5ʹ-methylcytosines.

Adult zebrafish aged 5-10 months were anesthetized by immersion in a 0.16% Tricaine solution (MESAB, across Organics). Using a sapphire blade, a small cut was made in the sclera of the left eye. 0.5 μL of 500μM ASO were intravitreally injected into the incision site. An equal volume of 500μM GFP ASO was used as a control, while another group was injected with an equal volume of 1X phosphate-buffered saline (PBS) as an injection control. Zebrafish were pretreated with PTB ASOs 1 day prior to NMDA damage.

### NMDA damage Model

24 hours post ASO injection (hpi), zebrafish were anesthetized once again using Tricaine. An additional incision was made on the sclera of the left eye, and 0.5μL of a 100 mM NMDA solution was intravitreally injected. 1X PBS injections were used as an injection control for NMDA damage.

### Immunohistochemical staining

Fluorescent staining was performed as previously described (Konar et al., 2024). Zebrafish treated only with ASO injections were collected 3 days post injection (3dpi), while zebrafish undergoing additional NMDA damage were collected 3 days after NMDA injections. Upon collection, eyes were fixed in 4% paraformaldehyde overnight at 4°C. Eyes were then incubated in a 5% sucrose solution with 4 changes and kept overnight in 30% sucrose at 4°C. After sucrose treatment, eyes were placed in a 2:1 Cryomatrix (Fisher Scientific) and 30% sucrose solution for two hours at 4°C and then fully embedded in Cryomatrix. Eyes were sectioned at 15μm thickness using a Leica cryostat and placed on charged Histobond slides (Fisher Scientific). Slides were dried on a heating block and stored at −80°C. Prior to IHC staining, slides were placed on a heating block and then rehydrated in 1X PBS for 30 minutes. Slides were then subjected to antigen retrieval before being allowed to cool to room temperature (RT) (Konar et al., 2024). Slides were then placed in a blocking solution (3% donkey serum, 0.1% Triton X-100 in 1X PBS) for 2 hours at room temperature (RT) then incubated at 4°C in primary antibody solution overnight. Primary antibodies are listed in **Table 1**. Afterwards, slides were incubated in secondary antibody solutions with To-Pro-3 for 2 hours at RT. Secondary antibodies included donkey anti-mouse 488 (1:500), donkey anti-rabbit 488 (1:500), donkey anti-mouse cy3 (1:500), and donkey anti-rabbit (1:500) (Jackson Immuno). Slides were then mounted with VectaShield Antifade Mounting Medium (Vector Labs).

**Table 1.**
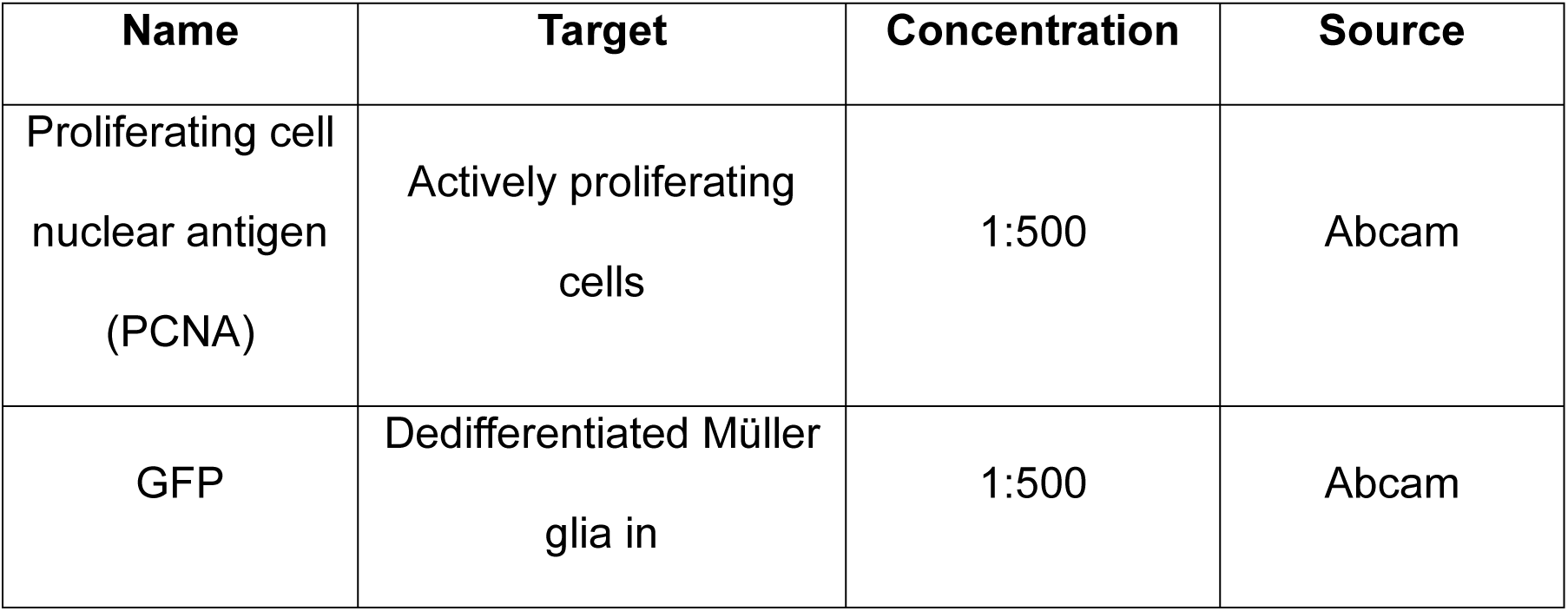
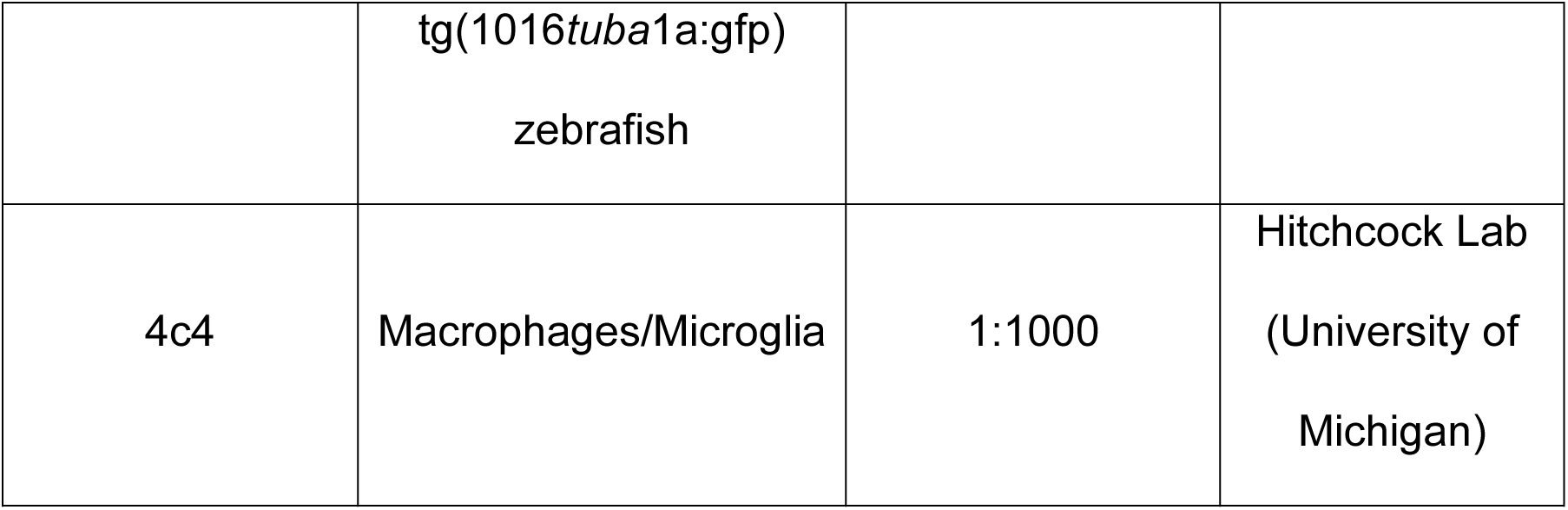
List of Antibodies Used For IHC Staining.

### Senescence-associated β-galactosidase staining

Slides were warmed and rehydrated in 1X PBS. Sections were then stained in accordance with Cell Signaling Technology kit instructions. Briefly, staining solution was warmed to 37°C then adjusted to a pH 6.0. Slides were incubated with staining solution overnight at 37°C. The staining solution was then washed off the slides, and slides were incubated in a 1:1000 dilution of To-Pro-3 (ThermoFisher) in 1X PBS for nuclear staining. Slides were then mounted using VectaShield antifade mounting medium (Vector Labs).

### Quantitative reverse transcription polymerase chain reaction (qRT-PCR)

qRT-PCR was performed on retinas collected 3 days post ASO injection as described (Kent et al., 2021). Retinas were placed in TriZol (Invitrogen) after dissection for total RNA purification. For cDNA synthesis, we used the Accuscript High-Fidelity 1^st^ Strand cDNA Synthesis kit. PCR amplification of cDNA was performed using the SYBR Green master mix (Bio-Rad Laboratories) with a Bio-Rad CFX96 real-time system. Amplification levels were normalized to 18S rRNA levels using the ΔΔCt method. 3 retinas were pooled per replicate. **Table 2** lists the primer sets:

**Table 2.**
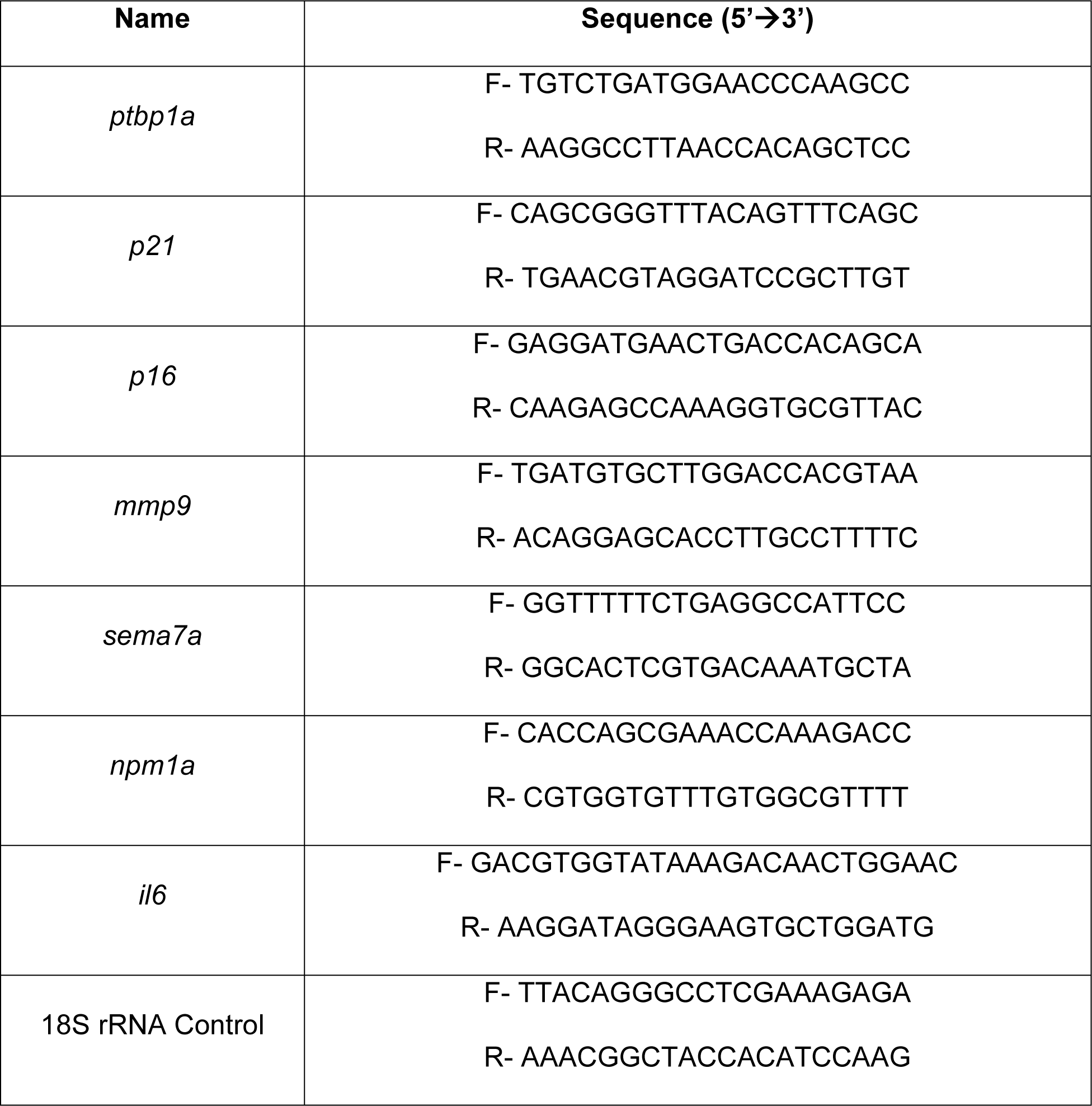
Primer sequences used for qRT/PCR.

### Imaging and image processing

IHC stains were imaged using a META Zeiss LSM Meta 510 confocal microscope in the Vanderbilt Center for Imaging Shared Resource (CISR) Core. SA-BGal and ToPRo were imaged on a NIKON AZ100M Widefield microscope. Confocal images were processed using ZEN Blue version 3.1, with additional analyses performed using ImageJ. AZ100M images were initially processed via NIS-Elements Viewer 5.21 and further analyzed with ImageJ.

### Quantification and statistical analysis

For experiments involving immunostaining and cell number quantification, sections were evaluated in an unbiased and blinded manner. Stained slides were evaluated in the central retina, and the dorsal and ventral regions were counted to include the entirety of the retinal region, spanning approximately ∼300-400 µm from the optic nerve in either direction. We excluded regions of extreme structural damage or near the ciliary marginal zone. Counts were obtained from at least two independent central retina sections and averaged for every eye section. We calculated significance using a two-way ANOVA with Turkey’s post hoc test for intergroup comparisons GraphPad Prism 10.0.1. Graphs demonstrate the average ± standard error of the mean. The number of fish used in each experiment are listed in the figure legends.

### Single-cell RNA sequencing analysis

Previously processed single-cell RNA sequencing (scRNAseq) datasets representing undamaged zebrafish retinas were used to generate a Seurat (v. 5.1.0) object (Hao et al., 2024; Hoang et al., 2020). To visualize localization of *ptbp1a* expression to specific cell types, normalized expression data was overlaid on Uniform Manifold Approximation and Projection (UMAP) coordinates for the undamaged retina samples and the plot1cell package (v. 0.0.0.9000) was used to summarize cluster-level expression data (Healy & McInnes, 2024; McInnes et al., 2018; Wu et al., 2022).

## Results

### Depletion of PTB Induces Proliferation

Antisense oligonucleotides (ASOs) have been used to deplete PTB after transfection into cultured cells, lentiviral shRNA delivery into mouse astrocytes, addition to human cortical organoid media, and by simple injection into mouse cerebrospinal fluid (Juliano et al., 2008; McDowall et al., 2024; Rinaldi & Wood, 2018; Sazani & Kole, 2003). We generated ASOs containing phosphorothioate linkages and 5’-methylcytosine modifications (Maimon et al., 2021) that target transcripts encoded by the two zebrafish genomic copies of PTB (PTBP1a and PTBP1b) (**Figure 1A)**. We tested the effects of intravitreal injection of these ASOs on DNA replication, as marked by expression of Proliferating Cell Nuclear Antigen (PCNA). Compared to control ASOs targeting GFP in otherwise undamaged retinas, we found that targeting *ptbp1a* led to significant increases in PCNA+ cells, whereas targeting *ptbp1b* had no effect (**Supplemental Fig. 1**). We therefore focused on *ptbp1a* and found that intravitreal injection of *ptbp1a* ASOs successfully reduced *ptbp1a* expression levels by ∼70% compared to PBS-injected controls (**Figure 1B**). Control ASOs targeting GFP did not cause significant reduction in *ptbp1a* expression.

**Figure 1.**
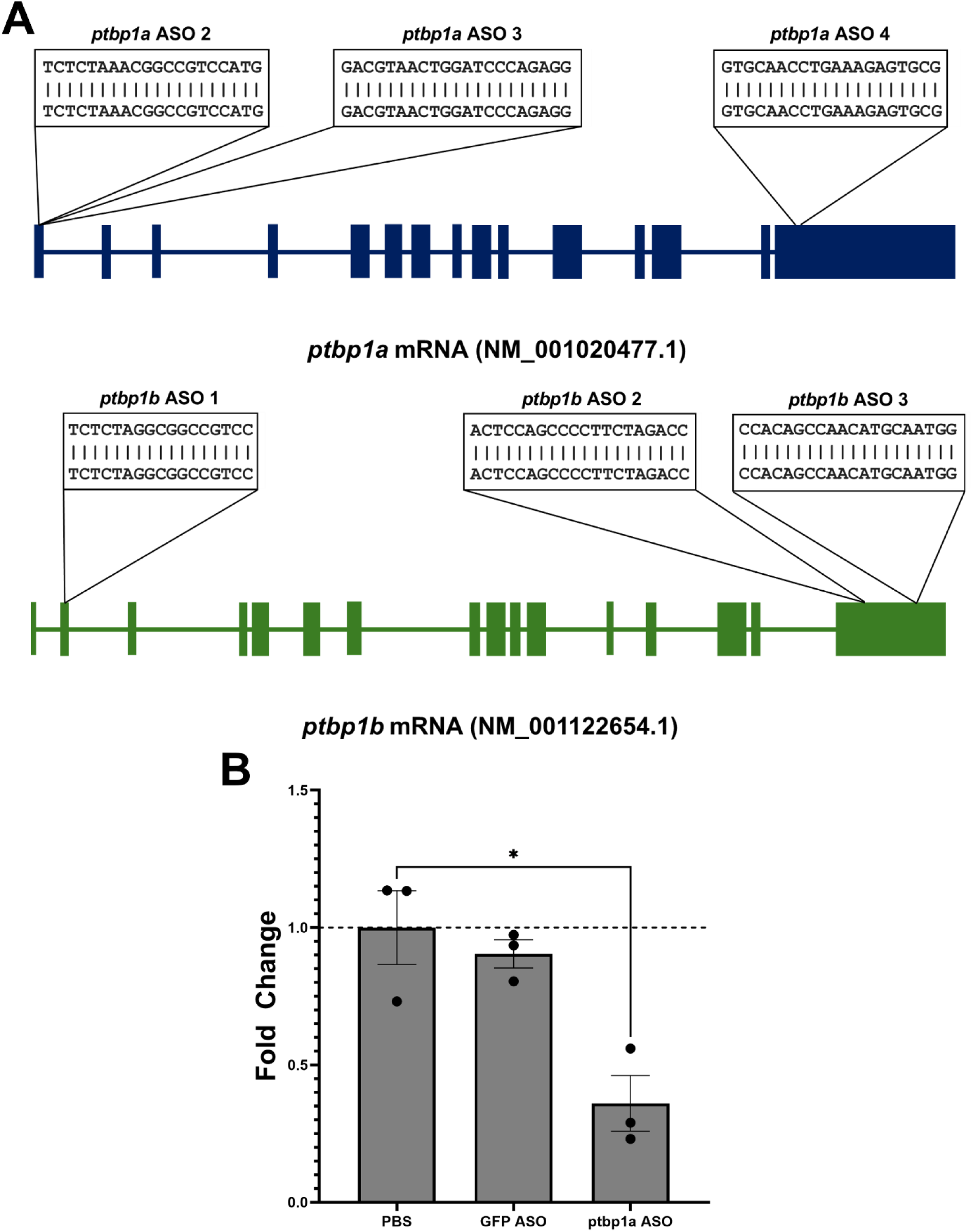
Targeting *ptbp1a* and *ptbp1b* with antisense oligonucleotides. (A) Zebrafish contain two copies of PTBP1 (a and b) with the indicated exon and intron structures. Antisense oligonucleotides (ASOs) were synthesized complementary to the boxed regions of *ptbp1a* and *ptbp1b.* (B) Adult zebrafish were injected with ASOs targeting *ptbp1a*. At 3 days post injection (dpi), retinas were dissected and RNA was purified for qRT/PCR of *ptbp1a* levels with fold-changes normalized to PBS injected control retinas. (n=3, * = p <0.05)

### MG-derived proliferation increases after *ptbp1a* knockdown

To determine whether the increase in proliferation observed after injection of ASOs targeting *ptbp1a* is a regeneration-associated response mediated by MG, we used *Tg[1016;tuba1a;GFP]* transgenic zebrafish. In this line, GFP marks dedifferentiated MG and MG-derived progenitor cells (Fausett & Goldman, 2006). We also tested how injection of *ptbp1a* ASOs would affect proliferation when combined with retinal damage by injection of NMDA, which causes significant neuronal death in the ganglion cell layer (Dvoriantchikova et al., 2023). We thus intravitreally injected *Tg[1016;tuba1a;GFP]* fish with *ptbp1a* ASOs followed by a second injection after 24hr. of either PBS or NMDA 24. Retinas were then collected at 3dpi and immunostained for GFP and PCNA. In control retinas in the absence of NMDA damage, no proliferation was observed after injection of either PBS or ASOs targeting GFP (**Figure 2A,B**). Injection of ASOs targeting *ptbp1a* in the absence of NMDA damage resulted in the detection of numerous PCNA+ cells in the Inner Nuclear Layer (INL), some of which co-localized with GFP. When NMDA damage was combined with injection of *ptbp1a* ASOs, we observed a striking increase in the number of PCNA+ cells in the INL and the majority of these cells co-localized and formed clusters with GFP+ cells (**Figure 2A,B**). The formation of MG-derived clusters of proliferating cells (GFP+/PCNA+) in the INL is characteristic of bona fide regeneration (Rajaram et al., 2014).

**Figure 2.**
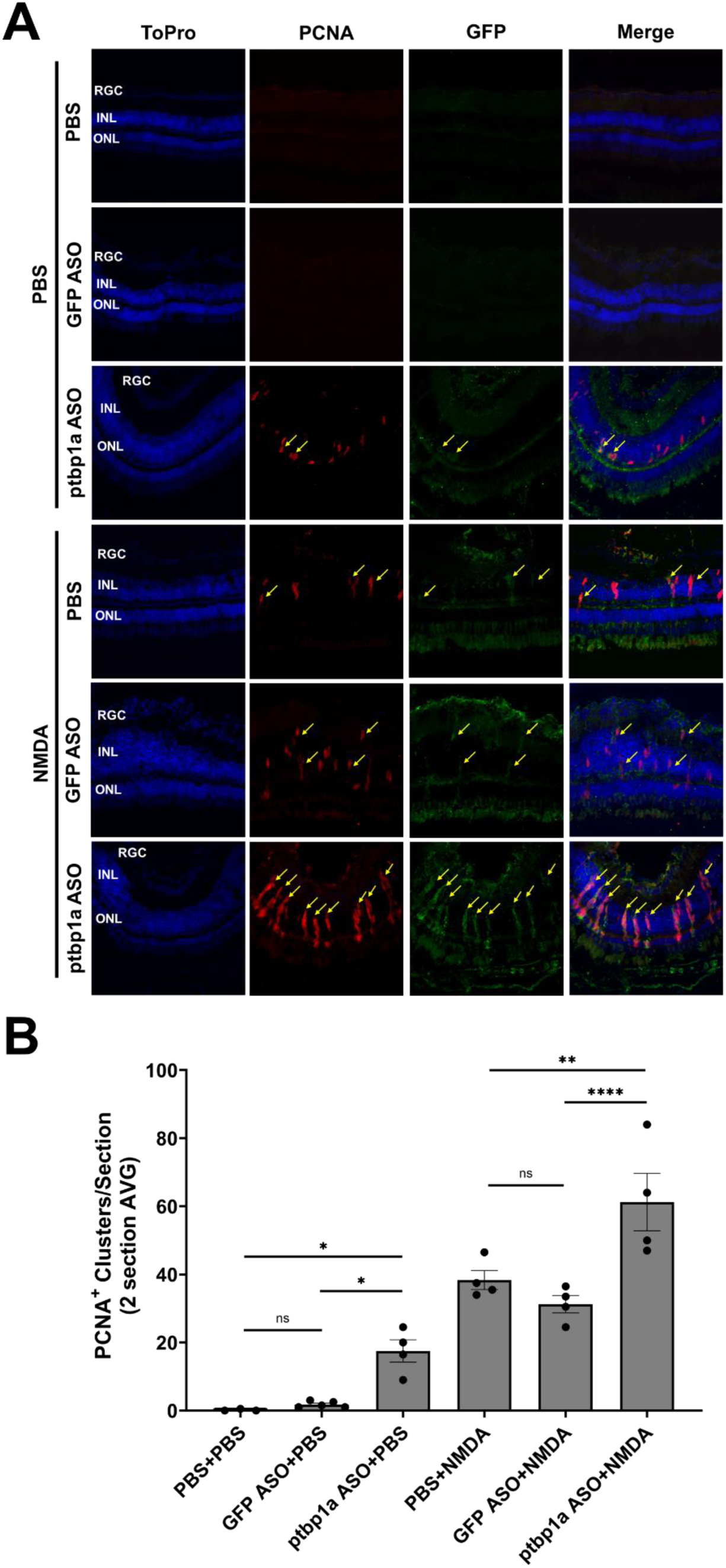
*ptbp1a* knockdown induces Müller glia-derived proliferation. Tg[1016;tuba1a;GFP] transgenic zebrafish were intravitreally injected with ASOs targeting either GFP or *ptbp1a* in the presence or absence of NMDA. (A) 24 hours after ASO or control PBS injections, a second injection of PBS or NMDA was administered followed by immunostaining at 3dpi for the nuclear marker ToPro, PCNA or GFP. (B) Quantification of the number of PCNA+/GFP+ clusters in the Inner Nuclear Layer. Co-localization of PCNA and GFP is indicated by arrows. (n=3-6, * = p <0.05, ** = p<0.01, *** = p<0.001, **** = p,0.0001)

### *ptbp1a* knockdown induces a senescent, pro-inflammatory response

One possible mechanistic explanation to resolve the controversy over the role of PTB during retina regeneration is that depletion of PTB activates regeneration through control of secreted factors that affect the local microenvironment and induce a MG response. Consistent with this, depletion of PTB has been shown to regulate the inflammatory secretome and induce cellular senescence (Georgilis et al., 2018; Hensel, Nicholas, Kimble, Nagpal, Omar, Tyburski, Jellison, Menoret, et al., 2022; Li et al., 2024; Yang et al., 2024). If reduced expression of PTB can induce or promote senescence in the retina, it would be consistent with our previous discovery of a key role for senescent cells during retina regeneration (Konar et al., 2024). To test this, we injected wildtype zebrafish retinas with ASOs targeting *ptbp1a* and quantified the impact on senescence markers and senescence gene expression. At 3dpi, we found that compared to ASOs targeting GFP, *ptbp1a* knockdown significantly increased the number of senescent cells as marked by expression of Senescence Associated β-galactosidase (SA-βgal^+^), a traditional marker of senescent cells (Dimri et al., 1995; Itahana et al., 2007) (**Figure 3A,B**). We also observed upregulation of two additional markers of senescence that regulate cell cycle arrest, *p16* and *p21*, via qRT/PCR (Dodig et al., 2019; Schmitt et al., 2002) (**Figure 3C-D**). These data are consistent with depletion of PTB inducing a senescent response (Li et al., 2024; Yang et al., 2024).

**Figure 3.**
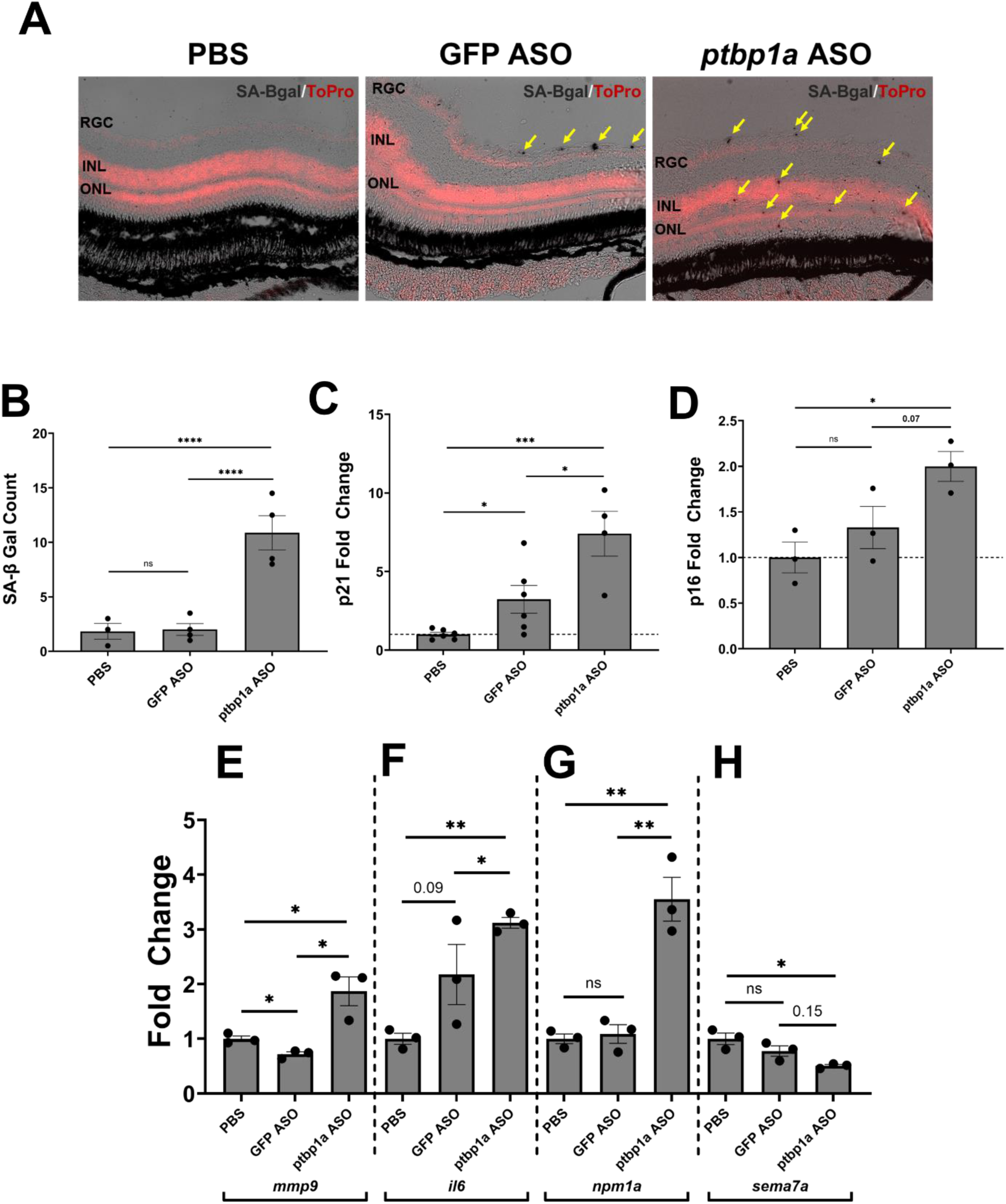
ptbp1a knockdown induces expression of senescence markers and SASP factors. Adult wild type zebrafish were intravitreally injected with PBS or ASOs targeting GFP or ptbp1a. (A) At 3dpi, sections were stained for the senescence marker SA-βgal^+^ (arrows). (B) Quantification of SA-βgal^+^ cells after injection of PBS or ASOs (C) Quantification of p21 levels after injection of PBS or ASO ASOs. (D) Quantification of p16 levels after injection of PBS or ASOs. (E-H) RNA was isolated from retinas at 3dpi after the indicated intravitreal injections and qRT/PCR was performed to quantify the relative levels (compared to control PBS injections) of the pro-inflammatory SASP factors mmp9, and IL-6, the pro-regenerative SASP factor npm1a, and semaphorin 7a, which negatively regulates the immune system. (n=3-6, * = p <0.05, ** = p<0.01, *** = p<0.001, **** = p,0.0001)

The appearance of senescent cells after depletion of *ptbp1a* predicts that SASP factors should also be upregulated. We thus isolated RNA at 3 dpi and performed qRT/PCR to analyze the expression of key SASP factors (Basisty et al., 2020). We observed a significant increase in the expression of the SASP factors matrix metalloproteinase 9 (*mmp9*) and interleukin-6 (*il6*) (**Figure 3E-F**). This is consistent with *ptbp1a* regulation of the pro-inflammatory SASP secretome (Georgilis et al., 2018). Additionally, we found that *ptbp1a* knockdown caused upregulation of the pro-regenerative factor nucleophosmin 1 (*npm1a*), which has been shown to regulate MG-derived progenitor production through *chx10/vsx2* (**Figure 3G**) (Basisty et al., 2020; Konar et al., 2025; Ouchi et al., 2012). Finally, we observed a decrease in the expression of semaphorin 7a (*sema7a*), which is a negative regulator of the immune system, further suggesting that *ptbp1a* knockdown influences inflammation and SASP expression (Czopik et al., 2006) (**Figure 3H**). Combined, the data support the finding that depletion of *ptbp1a* induces senescence and is consistent with a role for SASP factors in the activation of regeneration.

### *ptbp1a* expression is localized to glial and endothelial cells in the retina

We previously showed that the primary senescent cell type that is detectable after retina damage in zebrafish is derived from microglia/macrophages, as marked by immunostaining with 4c4 antibodies (Konar et al., 2024; Rovira et al., 2022). After intravitreal injection of *ptbp1a* ASOs, we observed an increase in the number of 4c4+ cells in the retina with a larger increase observed when combined with NMDA damage (**Figure 4A,B**). This is consistent with localization of *ptbp1a* expression in microglia, as determined by scRNAseq analyses (**Figure 4C, D**). Interestingly, *ptbp1a* expression in undamaged retinas was also detected in resting MG, activated MG, MG-derived progenitor cells, and vascular endothelial cells (**Figure 4E).** Analysis of scRNAseq data sets showed that these same cells express multiple SASP factors, one of which is PTB (Konar et al., 2025).

**Figure 4.**
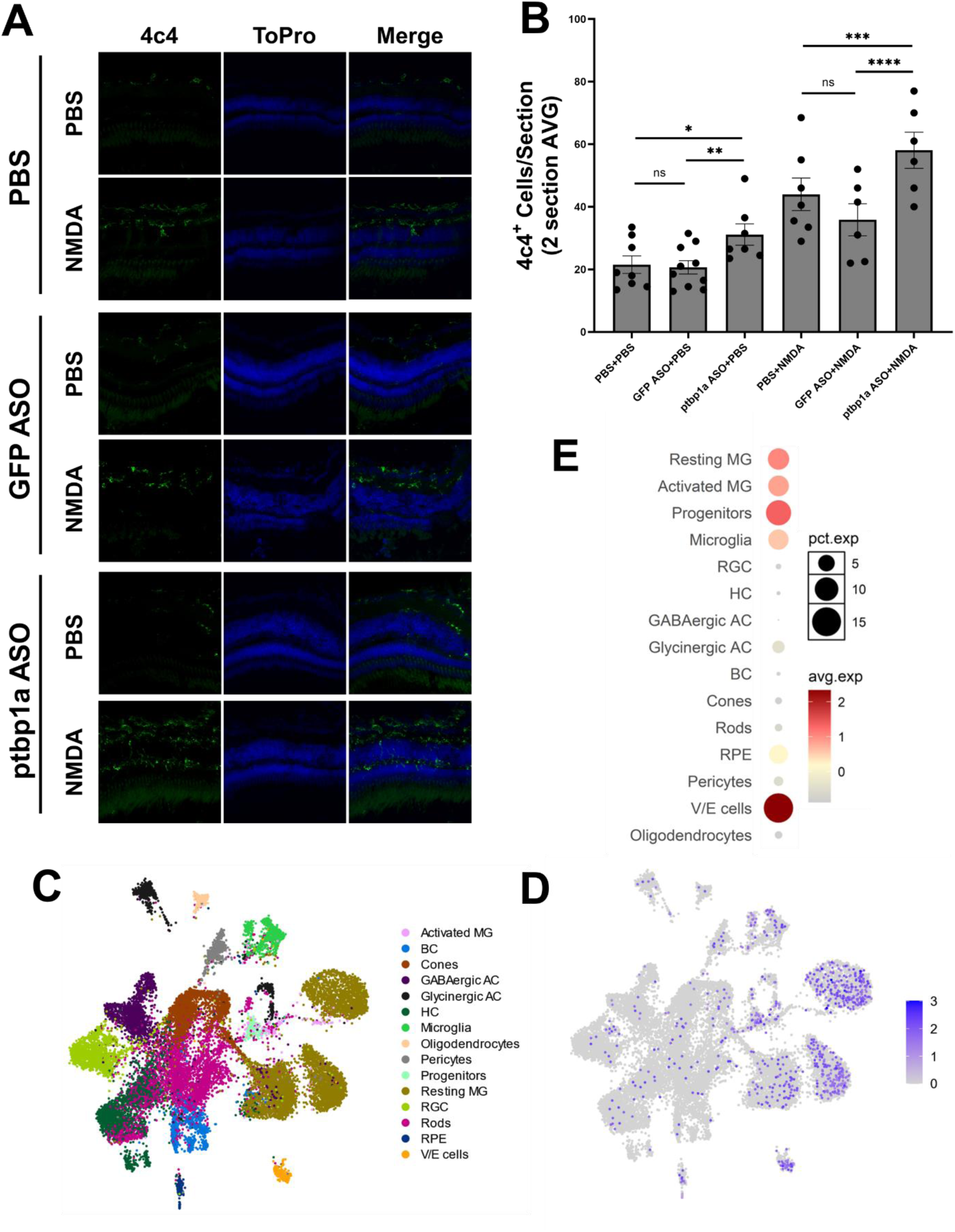
*ptbp1a* expression is localized to glial cells and causes increased immune cell numbers after knockdown. (A) Wild type zebrafish were intravitreally injected with either PBS or ASOs targeting GFP or *ptbp1a* in the presence or absence of NMDA damage. At 3pi, sections were immunostained with ToPro or 4c4 antibodies which recognize microglia/macrophages. (B) Quantification of 4c4+ cells under the indicated conditions. (C-D) UMAP visualization of cell types expressing *ptbp1a* in undamaged retinas. Cell types are indicated in the legend at the right. *ptbp1a* expression (blue) in the same cells is depicted in panel D. (E) Quantification of *ptbp1a* expression visualized in dot plots illustrating the percent of each cell type expressing *ptbp1a* (dot size) and average expression indicated by the red color gradient. (n=7-9, * = p <0.05, ** = p<0.01, *** = p<0.001, **** = p,0.0001)

## Discussion

Here, we show that knockdown of PTB (*ptbp1a*) in the zebrafish retina induces a senescent response with proliferation of MG-derived progenitor cells. It is known that inflammation must be properly modulated for retina regeneration (Bludau et al., 2024; Iribarne & Hyde, 2022b; Leach et al., 2021; Mitchell et al., 2018; Nagashima & Hitchcock, 2021; Zhou et al., 2022) and our discovery that depletion of *ptbp1a* leads to upregulation of pro-inflammatory SASP factors such as *mmp9* and *Il6* is consistent with an early requirement for inflammation. Strikingly, we find that depletion of PTB can induce MG-derived proliferation in undamaged retinas although the effects are much more robust in the presence of damage.

### PTB and Retina Regeneration

Both shRNA knockdown and overexpression of *miR-124* have shown that decreased levels of PTB can induce neuronal differentiation, including direct conversion of fibroblasts to neurons (Makeyev et al., 2007; Xue et al., 2013). Those studies prompted experiments to package either PTBP1-targeting shRNA vectors or CRISPR/CasRx into AAV vectors for delivery to the brain and retina and resulted in the surprising finding that knockdown of *ptbp1* could induce direct glia-to-neuron conversion (Qian et al., 2020; Zhou et al., 2020). It was later found that the vectors used to deplete *ptbp1* were not specific for MG and cell-lineage experiments and genetic knockout of PTB also did not support direct neural conversion (Hoang et al., 2023; Wang et al., 2021; Xie et al., 2022). Despite the controversy, multiple studies support the finding that knockdown of PTB induces gene expression changes by mechanisms that remain unclear but which are consistent with neuronal differentiation (Maimon et al., 2021; Makeyev et al., 2007; Xue et al., 2013). Genetic compensation or transcriptional adaptation might explain differences between the PTB knockout and knockdown studies (El-Brolosy & Stainier, 2017; Fu & Mobley, 2023; Rossi et al., 2015). This could be particularly relevant for PTB studies because neuronal induction and maturation are controlled by overlapping regulatory loops that involve *miR-124, miR-9*, and the REST and BRN2 transcription complexes (Hu et al., 2018). Any of the individual factors within these loops could conceivably be transcriptionally altered by knockouts (and knockdowns) of PTB (Fu & Mobley, 2023). Another non-mutually exclusive explanation could be that depletion of PTB from immune-derived cell populations, a subset of MG, or vascular endothelial cells induces a senescent response and that subsequent changes in the SASP secretome provide pro-inflammatory signals within the microenvironment that initiate a proliferative, MG-derived regenerative response. This is consistent with a role for depletion of PTB in inducing senescence and inflammation (Georgilis et al., 2018; Hensel, Nicholas, Kimble, Nagpal, Omar, Tyburski, Jellison, Ménoret, et al., 2022; Li et al., 2024; Yang et al., 2024).

### PTB, Senescence, and Inflammation

One major facet of PTB functionality is that it plays a role in both senescence regulation and inflammatory state modulation (Georgilis et al., 2018; Hensel, Nicholas, Kimble, Nagpal, Omar, Tyburski, Jellison, Ménoret, et al., 2022; Li et al., 2024; Yang et al., 2024). Zebrafish are unique in that they can modulate their inflammatory status from highly pro-inflammatory at the beginning of regeneration, to a more anti-inflammatory and pro-regenerative environment during later stages of regeneration (Bludau et al., 2024; Iribarne & Hyde, 2022b; Nagashima & Hitchcock, 2021; Zhou et al., 2022). This is a consistent with the transient detection and clearance of senescent cells as regeneration proceeds (Konar et al., 2024). Knockdown of PTB in zebrafish induced an increase in immune cell numbers (4c4+ cells) (**Figure 4A,B**) and also upregulated numerous genes associated with the pro-inflammatory state (**Figure 3E-F**). This suggests that PTB is playing a role in modulating the inflammatory state of the retina. The pro-inflammatory stage is often correlated with MG reactivity, consistent with the observed increase in MG-derived proliferation (**Figure 2A,B**) (Bludau et al., 2024; García-García et al., 2024; Nelson et al., 2013). Together, our data support a model whereby depletion of *ptbp1* induces a change in SASP factor expression in the retina and that such microenvironmental changes can initiate a MG-derived proliferative response.

## Acknowledgements

The authors would like to thank Jack Hollander for his assistance in maintaining the zebrafish lines used in this manuscript.

## Author Contributions

G.K.- Conceptualization, data curation, formal analysis, methodology, writing- original draft, writing- review and editing. A.L.- data curation, formal analysis, methodology, writing- original draft, writing- review and editing. K.V.- data curation, formal analysis, software, writing- review and editing. T.N.- data curation. Z.F.- conceptualization, data curation. J.P.- conceptualization, funding acquisition, project administration, resources, supervision, writing- review and editing.

