## Supplemental Figure 1 for "Depletion of Polypyrimidine tract binding protein 1 (*ptbp1*) activates Müller glia-derived proliferation during zebrafish retina regeneration via modulation of the senescence secretome"

**Konar et al Supplemental Figure 1**

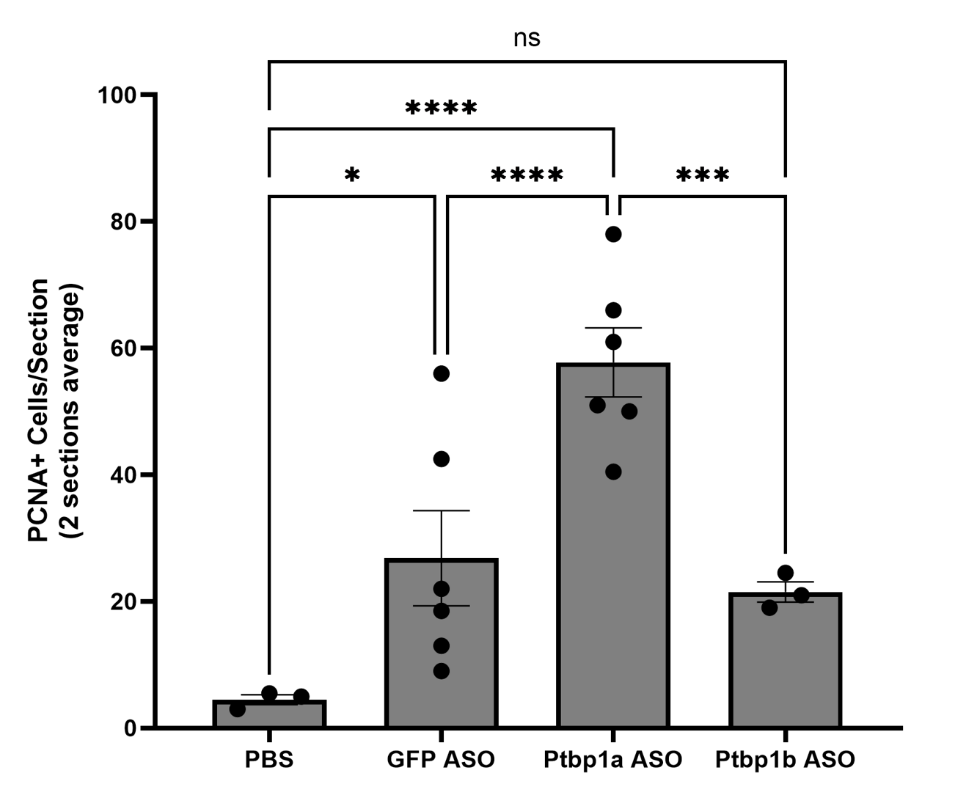

**Supplemental Figure 1: *ptbp1a* is the active isoform of PTB for proliferation control.** Wild type adult zebrafish were intravitreally injected with either PBS or the indicated ASOs. At 3dpi, retinal sections were immunostained for PCNA.
